# Tolerance of an acute warming challenge declines with body mass in Nile tilapia: evidence of a link to capacity for oxygen uptake

**DOI:** 10.1101/2020.12.03.409870

**Authors:** F.R. Blasco, E.W. Taylor, C.A.C. Leite, D.A. Monteiro, F.T. Rantin, D.J. McKenzie

## Abstract

It is proposed that larger individuals within fish species may be more sensitive to global warming, due to limitations in their capacity to provide oxygen for aerobic metabolic activities. This could affect size distributions of populations in a warmer world but evidence is lacking. In Nile tilapia *Oreochromis niloticus* (n=18, mass range 21-313g), capacity to provide oxygen for aerobic activities (aerobic scope) was independent of mass at acclimation temperature (26°C). Tolerance of acute warming, however, declined significantly with mass when evaluated as critical temperature for fatigue from aerobic swimming (CT_swim_). The CT_swim_ protocol challenges a fish to meet the oxygen demands of constant intense aerobic exercise while their demands for basal metabolism are accelerated by incremental warming, culminating in fatigue. CT_swim_ elicited pronounced increases in oxygen uptake but maximum rates achieved prior to fatigue declined very significantly with mass. Mass-related variation in CT_swim_ and maximum oxygen uptake rates were positively correlated, which may indicate a causal relationship. When faced with acute thermal stress, larger fishes within populations may become constrained in their ability to swim at lower temperatures than smaller con-specifics. This could affect survival and fitness of larger fish in a world with more frequent and extreme heatwaves, with consequences for population productivity.

## Introduction

There is evidence that ongoing global warming is associated with a progressive decline in final asymptotic body size in many fish species (Audzijonyte et al., 2020; Baudron et al., 2014; Daufresne et al., 2009; Gardner et al., 2011). This phenomenon may have a physiological mechanism, understanding this could improve the ability to predict effects of future warming (Audzijonyte et al., 2019; Lefevre et al., 2017; Lefevre et al., 2018). There has been major focus upon respiratory gas-exchange in relation to size and temperature in fishes, because water is relatively poor in oxygen and meeting requirements for aerobic metabolism can be challenging (Audzijonyte et al., 2019; Cheung et al., 2011; Lefevre et al., 2017).

The Gill Oxygen Limitation (GOL) model proposes a physiological mechanism to explain declining adult fish sizes. The model posits that, as fishes grow, gill respiratory surface area declines in relation to body volume, due to surface to volume relationships of spherical bodies. As fishes grow, therefore, their capacity to meet the oxygen demands of their body volume, and hence body mass, would decrease progressively. At a certain body size, the gills would only be able to meet the oxygen requirements of basal metabolism. At that size, the fish would have no aerobic metabolic scope (AS) to provide oxygen for aerobic activities, including anabolism, so growth would no longer be possible (Cheung et al., 2011; Pauly, 1981). Fishes are ectotherms so, when warmed, their metabolic rate and associated oxygen demand increase (Fry, 1957; Fry, 1971). In the GOL model, increased basal metabolic demands would cause the limitation in gill oxygen uptake capacity to occur at smaller maximum sizes. The model has, consequently, been used to project widespread global declines in fish size due to environmental warming (Cheung et al., 2011; Cheung et al., 2012; Pauly and Cheung, 2017). Numerous major physiological precepts of the GOL model are not, however, supported by current knowledge or data (Audzijonyte et al., 2019; Killen et al., 2016; Lefevre et al., 2017).

Tolerance of acute warming does, nonetheless, decline with mass in some fish species (Leiva et al., 2019; McKenzie et al., 2020). Although some studies have considered whether size-dependent tolerance might be linked to capacity for oxygen uptake (Christensen et al., 2020; Messmer et al., 2017), this remains to be tested explicitly. This study represents an experimental test of the GOL model and the potential effects of body mass on tolerance of acute warming in a teleost fish. The Nile tilapia *Oreochromis niloticus* is a relatively eurythermal teleost (natural thermal range 14-33°C) that provides important fisheries in tropical and sub-tropical countries across the globe (De Silva et al., 2004; Schofield et al., 2011). We performed a series of tests on tilapia that ranged over one order of magnitude in body mass, to (1) how tolerance of acute warming challenges varied with mass and (2) whether this could be related to variation in capacity for oxygen uptake with mass.

In order better to interpret any effects of mass on thermal tolerance, we first measured key traits of respiratory metabolism and performance at acclimation temperature (26°C), by swimming respirometry. We then investigated effects of mass on tolerance of warming using two thresholds. The critical thermal maximum (CT_max_) protocol is the standard method to evaluate tolerance, fish are warmed incrementally and loss of equilibrium (LOE) is the tolerance endpoint (Beitinger and Lutterschmidt, 2011). The critical temperature for aerobic swimming (CT_swim_) protocol warms fish incrementally while they perform intense steady aerobic exercise in a swimming respirometer, with fatigue as tolerance endpoint (Blasco et al., 2020b). The CT_swim_ evaluates capacity for oxygen uptake because it challenges a fish to meet the combined oxygen demands of aerobic exercise (constant metabolic load) plus progressive warming (incremental metabolic load). Warming causes a progressive and profound increase in oxygen uptake up to a maximum rate, where the fish transitions to unsustainable anaerobic swimming that presages imminent fatigue (Blasco et al., 2020b). We therefore investigated whether variation in CT_swim_ with mass was correlated with maximum oxygen uptake at fatigue, as evidence of a causal relationship between the variables.

## Material and Methods

Experimental animals came from populations of Nile tilapia of different size/age classes, reared in recirculating biofiltered water at 25±1°C in multiple outdoor tanks (vol. 1000L) at the Department of Physiological Sciences, UFSCar, São Carlos (SP), Brazil. Individuals (n=18, mass range 20.6g to 313.0g) were selected in three sequential groups of six relatively size-matched individuals, tagged for identification (PIT under benzocaine anaesthesia) then recovered for at least 96h in an indoor tank (vol. 100L) in the same biofiltered water system and with routine feeding. Animals were fasted 24h prior to experimentation, and weighed prior to overnight recovery in an experimental apparatus. Animals did not change mass between sequential tests. Experimental protocols were approved by CEUA/UFSCAR, number CEUA 3927151016.

Metabolic and performance phenotype at acclimation temperature was measured with a Steffensen-type swim-tunnel (volume 13.4L) supplied with vigorously aerated, biofiltered water at 26±0.1°C, as described in (Blasco et al., 2020b). Briefly, fish were placed in the tunnel and left overnight at a low swimming speed equivalent to 1 bodylength·s^−1^ (BL·s^−1^, corrected for solid blocking effect) (Blasco et al., 2020b). The following morning swimming speed was increased each 30 min in steps of either 1 BL (mass range 20.6 to 86.5 g, n=12) or 0.5 BL (204 to 313 g, n=6), up to either 5 or 2.5 BL·s^−1^, respectively (Blasco et al., 2020a). All fish engaged steady aerobic body-caudal swimming at all these speeds (Blasco et al., 2020b). Measurements of oxygen uptake (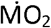, mmol·kg^−1^·h^−1^) were made by stopped-flow respirometry at each speed, to derive standard metabolic rate (SMR) (Chabot et al., 2016) and active metabolic rate AMR (Norin and Clark, 2016), and calculate AS as AMR-SMR (Blasco et al., 2020b). Subsequently, the swimming speed was increased by 0.1 BL every 10s to identify gait transition speed (U_GT_), where fish first engaged an unsteady anaerobic ‘burst-and coast’ gait, which then increased in intensity until maximum swimming speed (U_max_) and fatigue (Blasco et al., 2020b; Marras et al., 2013). Speed was then immediately reduced to 1 BL·s^−1^ and, after 30 min, fish were returned to a second holding tank for at least 96h under normal rearing conditions, prior to testing thermal tolerance (50% tested for CT_swim_ or CT_max_ first).

CT_max_ was measured on groups of tilapia that had acclimated overnight in a 68L tank containing vigorously aerated water at 26°C. The following morning, water was warmed 1 °C every 30 min until loss of equilibrium (LOE) (Blasco et al., 2020b). Individual CT_max_ was then recorded as the highest temperature step fully completed plus the proportion of the last step that the fish endured prior to LOE (25). Immediately upon LOE, fish were recovered at 26°C for at least 30 min, in a 68L tank of aerated water, then returned to a holding tank for at least 96h prior to further experimentation.

For CT_swim_ each individual recovered overnight in the swim-tunnel at 1 BL·s^−1^ then, the next day, speed was increased over 30 min until 85% of its own U_GT_ (Blasco et al., 2020b). After 30 min at that speed, temperature was increased by 1°C every 30 min until fish fatigued, resting against the rear screen (Blasco et al., 2020b). The CT_swim_ was calculated as for CT_max_ but using fatigue as endpoint (Blasco et al., 2020b). The fish was immediately placed in a recovery tank at 26°C for 30 min, then returned to holding tank for at least 96h prior to further experimentation. Measurements of 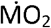 were made over the last 20 min at each temperature, the highest rate achieved was 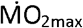 (Blasco et al., 2020b). Nile tilapia acclimated to 26°C can swim at a steady elevated 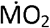 without fatiguing for at least 9h at 85% of U_GT_, exceeding the duration of the CT_swim_ (Blasco et al., 2020b).

Statistics were performed with SigmaPlot 11. Relationships with body mass, of metabolic, performance and tolerance variables, were assessed by least squares regression of log-transformed data. Correlations between variables (CT_max_ vs CT_swim_; CT_swim_ vs 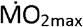) were assessed by Pearson product-moment. Mean CT_max_ and CT_swim_ were compared by paired T-test. P<0.05 was the limit for statistical significance.

## Results

At 26°C, log mass-specific SMR and AMR both showed a significant negative relationship with log body mass (Table 1, Figure 1A). Log mass-specific AS did not change with mass (Table 1, Figure 1A). Note that the negative slopes of these log:log relationships for the metabolic variables (Table 1, Figure 1A) were the reciprocals of the positive log:log slopes of mass independent scaling exponents for each variable (Fig. S1, Electronic Supplementary Material). Log U_GT_ and U_max_ both fell significantly with mass but the difference between them (U_max_-U_GT_), a potential indicator of anaerobic swimming capacity, was independent of mass (Table 1).

**Figure 1.**
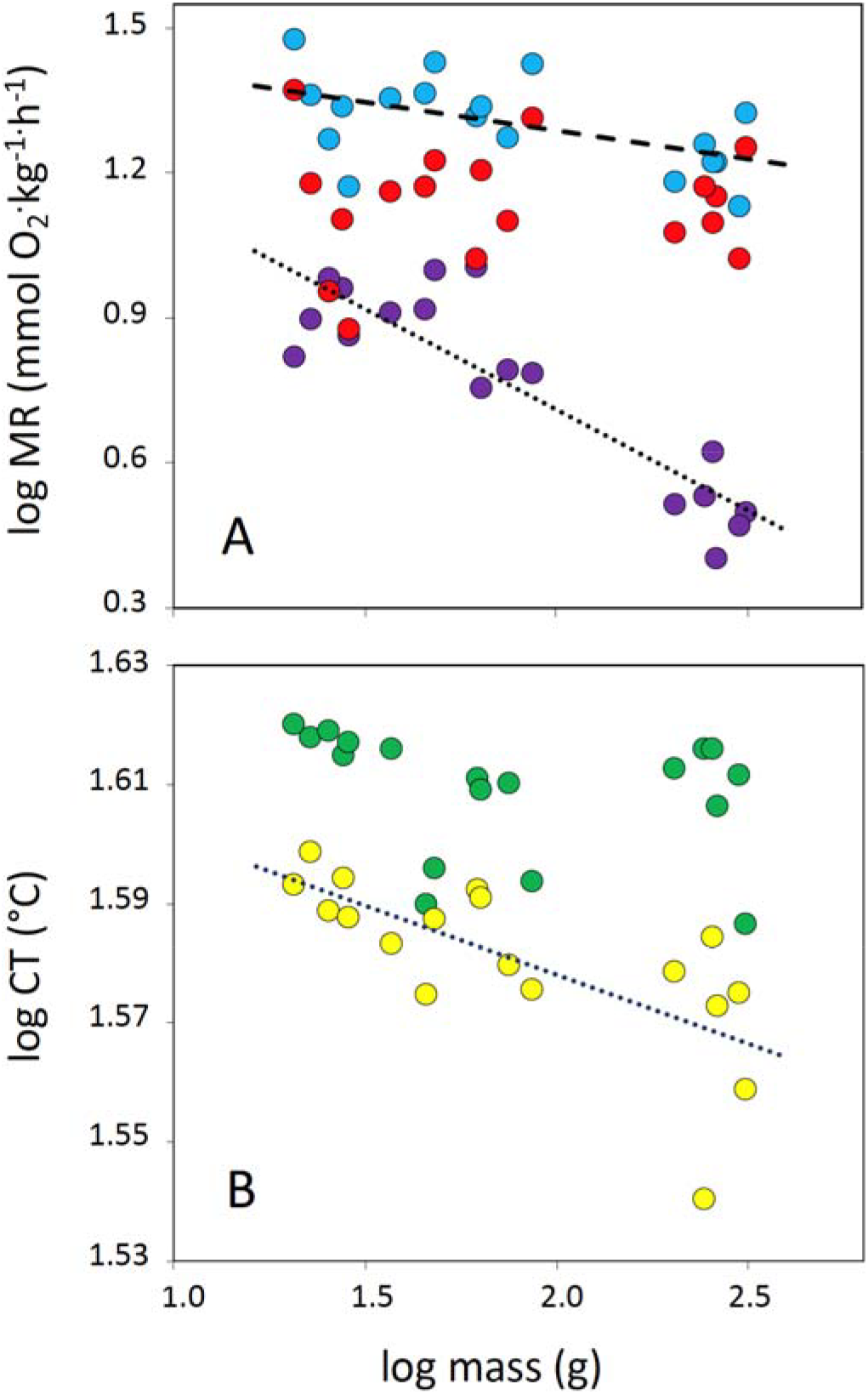
Least squares regression of the log:log relationship between (A) body mass and mass-specific metabolic rate (MR) variables, standard metabolic rate (purple symbols, dotted line), active metabolic rate (blue symbols, dashed line) and aerobic metabolic scope (red symbols); and (B) body mass and two critical temperatures (CT) for tolerance, critical thermal maximum (green symbols) and critical temperature for fatigue from swimming (yellow symbols, dotted line). Data are for n = 18 Nile tilapia, mass range 21 to 313g, acclimated to 26°C. Regression lines are for variables where relationship is significant, lines denote best fit, see Table 1.

As fish were warmed during CT_max_ they first exhibited erratic behaviour, rolling sideways, followed by complete LOE. During CT_swim_, fish swam using a steady aerobic gait up to a certain temperature, beyond which they progressively engaged unsteady burst and coast anaerobic swimming that led to fatigue. No fish lost equilibrium during CT_swim_; all fish from both protocols recovered normal swimming behaviour within 10 min at 26°C. The two thermal thresholds were not correlated (r=0.371, P=0.129) but overall mean (± SEM) CT_max_, at 40.7±23°C, was significantly higher than CT_swim_, at 38.1±0.29°C.

Log CT_max_ had no relationship with log body mass whereas log CT_swim_ declined significantly with mass (Figure 1B, Table 1). The decline in CT_swim_ with mass was associated with a highly significant negative log:log relationship between mass-specific 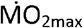 and mass (Figure 2A, Table 1, see Figure S1 for the reciprocal allometric scaling exponent). The 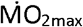 always occurred at the penultimate or ultimate temperature of the CT_swim_ except for one individual (mass 204g, see data in (Blasco et al., 2020a)). There was a significant positive correlation between log 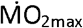 and log CT_swim_ (Figure 2B).

**Figure 2.**
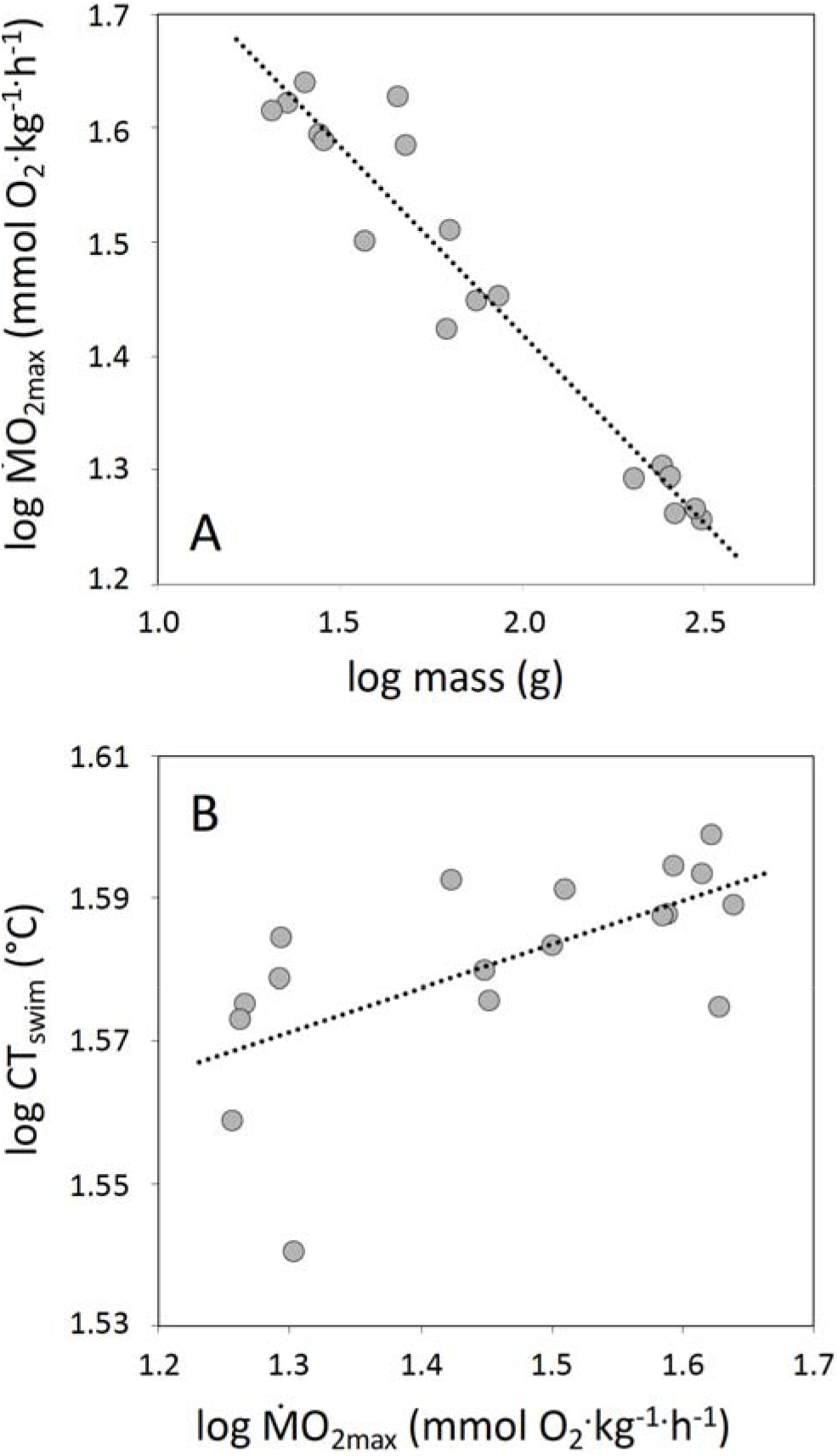
(A) Least squares regression of the log:log relationship between body mass and mass-specific maximum rate of oxygen uptake achieved during CT_swim_ (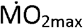, see Table 1 for regression information) and (B) Pearson correlation between critical temperature for fatigue from swimming (CT_swim_) and 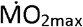, dotted line is described by log CT_swim_ = (log 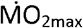) x 0.061 + 1.491 (r = 0.637, P = 0.004). Data are for n = 18 Nile tilapia, mass range 21 to 313g, acclimated to 26 °C.

## Discussion

The results demonstrate that the ability to perform sustained aerobic exercise, while being warmed (CT_swim_), declines significantly with body mass in a teleost fish. This decline was significantly correlated with a decline in mass-specific 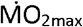 with mass, suggesting a causal relationship. This decline in 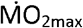 with mass during the CT_swim_ did not reflect differences in AS at acclimation temperature; AS was independent of mass. This latter finding is in direct opposition to the core precept of the GOL model, which predicts that AS will decline with increasing size (Cheung et al., 2011; Pauly, 1981). Nonetheless, there were declines in mass-specific SMR and AMR with mass at acclimation temperature, which were a direct reflection of allometric mass scaling relationships. These latter relationships occur in all fish species but their underlying mechanisms remain a matter of debate (Glazier, 2020; Killen et al., 2010; Killen et al., 2016).

The finding that CT_max_ did not decline with increasing body mass is consistent with a previous study on Nile tilapia (Recsetar et al., 2012). In studies that have investigated whether CT_max_ depends upon size in fishes, some report no effect but others report a significant decline with size (McKenzie et al., 2020). The physiological mechanisms underlying LOE at CT_max_ in fishes are unknown but it is notable that, when there is a dependence upon size in post-larval fishes, it is consistently negative (McKenzie et al., 2020). Given the absence of any dependence of CT_max_ upon mass, it was interesting to find such a strong negative dependence of CT_swim_.

It is intriguing that 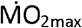 declined so profoundly with body mass during CT_swim_, if we assume that it represents the maximum capacity for oxygen uptake during acute warming (Blasco et al., 2020b). The significant correlation of 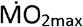 with CT_swim_ is evidence that capacity for oxygen uptake is a mechanism underlying size-dependence of CT_swim_. This is worthy of further investigation. The fact that size-related variation in 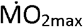 only accounted for about 60% of the variation in CT_swim_ indicates, however, that other mechanisms must also contribute to fatigue. The tilapia engaged unsustainable anaerobic swimming prior to fatiguing (Blasco et al., 2020b) and capacity for anaerobic exercise varies among individuals in fish species (Marras et al., 2010; Marras et al., 2013). At acclimation temperature, both U_GT_ and U_max_ fell significantly against mass, because mass was related to length in the tilapia (Blasco et al., 2020a) and relative swimming performance declines with length in most fish species (Bainbridge, 1958; Beamish, 1978). If the difference between them (U_max_-U_GT_) is considered an indicator of anaerobic metabolic capacity in fishes (Marras et al., 2013), this was independent of mass.

The decline in 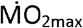 with mass presumably reflects, in some degree, the size-related constraints on mass-specific oxygen uptake capacity, AMR, at acclimation temperature. Such constraints were rendered very visible by the profound challenge to oxygen uptake that the CT_swim_ protocol imposed. Interestingly, the scaling exponent (1-*b*) for the decline in log mass-specific 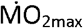 with log mass, of −0.33, is exactly the exponent that the GOL model posits for the decline in capacity for oxygen uptake as a function of mass in fishes, based upon surface to volume geometric constraints (Cheung et al., 2011; Cheung et al., 2012; Glazier, 2020; Pauly, 1981). In the tilapia, a regression coefficient of 0.92 on 18 fish indicates that estimated mass exponents are unlikely to be spurious. One constraint revealed by CT_swim_ may be gill functional respiratory surface area, which declines with mass in Nile tilapia with an exponent of-0.30 (Kisia and Hughes, 1992). This decline in gill surface area with mass may be linked to the decline in mass-specific SMR, whereby basal metabolic costs decline and the fish gills are then ‘tailored’ to maintain a constant AS at any given mass, as observed in this study. Maintaining a greater gill respiratory surface area than is needed for AS, at any given acclimation temperature, could have costs in terms of, for example, regulating water and ion balance (Lefevre et al., 2017; Taylor, 1998). There may be other surface to volume constraints that emerge as fishes grow and this is an interesting topic for further study, including on other fish species.

Extreme warming events are predicted to increase in frequency and severity in aquatic habitats worldwide due to accelerating global climate change (Field et al., 2012; Frölicher et al., 2018; Stillman, 2019). Swimming is an essential activity for most fishes, including Nile tilapia, so there are clear ecological implications to the fact that, if their populations experience an acute warming event, the ability of larger fishes to swim aerobically would be constrained at lower temperatures than that of smaller conspecifics. The law of averages would predict that larger fishes are more likely to encounter temperatures sufficiently warm to limit their performance (Field et al., 2012; Stillman, 2019). Therefore, stochastic extreme warming events could threaten fitness and survival of fishes in a size-dependent manner, including by limiting the ability of larger individuals to seek thermal refuges (Habary et al., 2017; Stillman, 2019). If fitness and survival of larger animals is compromised more frequently, this could have important consequences for Nile tilapia fisheries, an important source of dietary protein in many countries (De Silva et al., 2004).

Overall, the data indicate that global warming could indeed ‘favour the small’ for fishes (Daufresne et al., 2009). Further work is required to understand if this size-dependence in temperature tolerance might be linked to ongoing declines in maximum body size of fishes globally (Audzijonyte et al., 2019; Baudron et al., 2014; Daufresne et al., 2009). The rate of global warming is too fast for adaptive evolutionary responses by most fish species, so survival and resilience will depend upon the options for migration, the prevailing plasticity in tolerance, and the existence of tolerant genotypes (Bell, 2013; Gunderson and Stillman, 2015; Habary et al., 2017; Stillman, 2019). The current data indicate that capacity for migration is more likely to be impaired in larger fish, and that smaller animals are more tolerant. The potential downstream consequences, for future populations, may indeed be a decline in maximum body size.

## Acknowledgements

We are grateful to Cesar Polettini, of Piscicultura Polettini, for donating the tilapia.

## Funding Statement

FRB was supported by a doctoral bursary from Coordenação de Aperfeiçoamento de Pessoal de Nível Superior (CAPES).

## Notes

### Competing Interest Statement

The authors have declared no competing interest.

