## Supplementary material for "Tolerance of an acute warming challenge declines with body mass in Nile tilapia: evidence of a link to capacity for oxygen uptake": Figure S1

**Figure S1.** FR Blasco, EW Taylor, CAC Leite, DA Monteiro, FT Rantin, DJ McKenzie, Tolerance of an acute warming challenge declines with body mass in Nile tilapia.

Regressions of log mass-independent metabolic rate ($mmolO_2 \bullet h^{-1}$) against log mass (g) for n = 18 Nile tilapia, for standard metabolic rate (SMR); active metabolic rate (AMR); aerobic scope (AS), and maximum oxygen uptake rate achieved during the critical temperature for fatigue from aerobic swimming ($CT_{swim}$) protocol ($\dot{M}O_{2max}$). SMR = 0.581 x mass - 1.451 ($R^2 = 0.873$); AMR = 0.881 x mass - 1.474 ($R^2 – 0.958$); AS = 1.006 x mass -1.878 ($R^2 = 0.927$), and $\dot{M}O_{2max}$ = 0.671 x mass - 0.923 ($R^2 = 0.983$). All significant at $P < 1 \times 10^{-7}$.

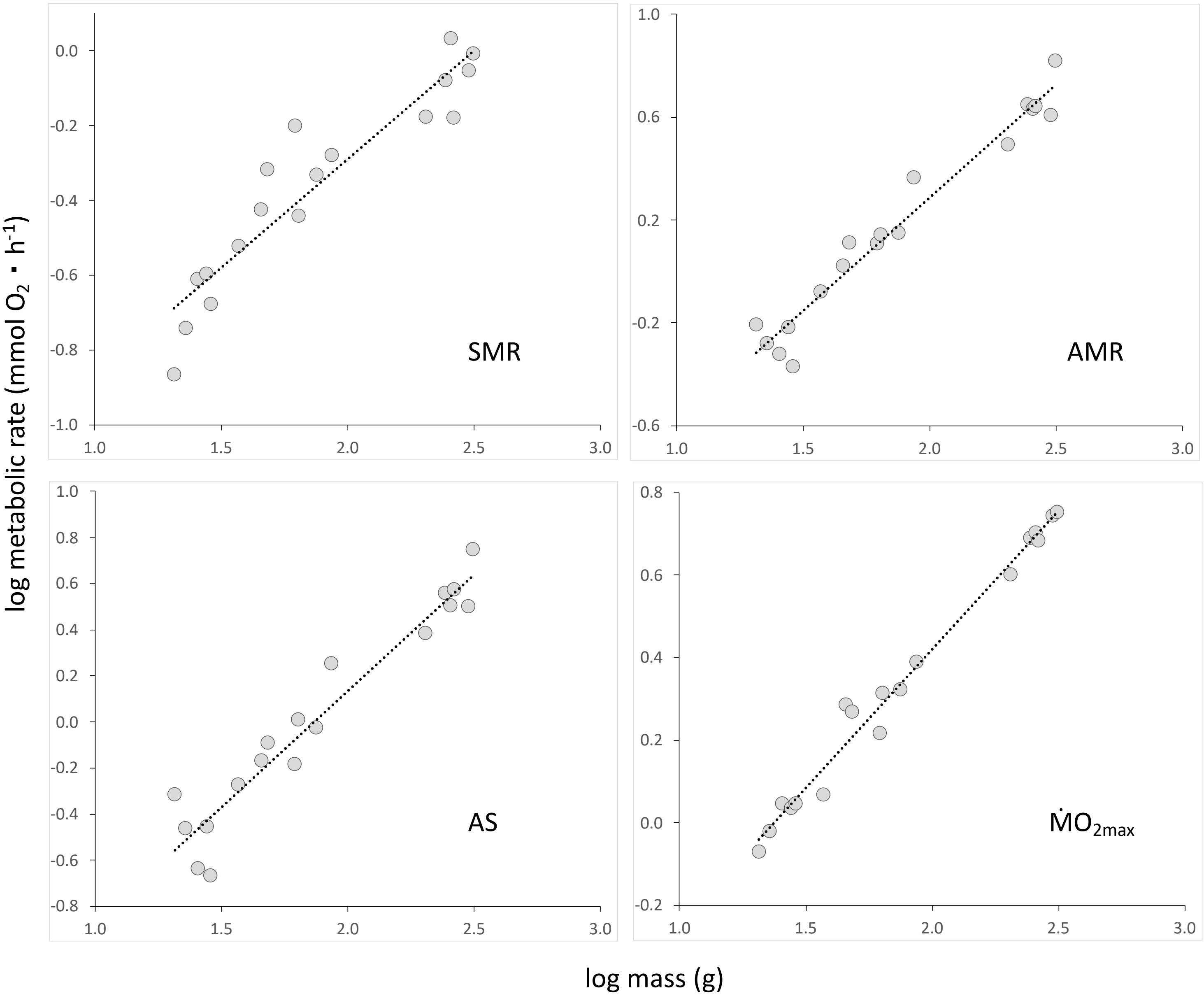
